# *CellSium* – Versatile Cell Simulator for Microcolony Ground Truth Generation

**DOI:** 10.1101/2022.03.24.485611

**Authors:** Christian Carsten Sachs, Karina Ruzaeva, Johannes Seiffarth, Wolfgang Wiechert, Benjamin Berkels, Katharina Nöh

**Author notes:** Equal contribution (Karina Ruzaeva, Johannes Seiffarth).

## Abstract

**Summary:** To train deep learning based segmentation models, large ground truth data sets are needed. To address this need in microfluidic live-cell imaging, we present *CellSium*, a flexibly configurable cell simulator built to synthesize realistic image sequences of bacterial microcolonies growing in monolayers. We illustrate that the simulated images are suitable for training neural networks. Synthetic time-lapse videos with and without fluorescence, using programmable cell growth models, and simulation-ready 3D colony geometries for computational fluid dynamics (CFD) are also supported.

**Availability and Implementation:** *CellSium* is free and open source software under the BSD license, implemented in Python, available at https://github.com/modsim/cellsium (DOI: 10.5281/zenodo.6193033), along with documentation, usage examples and Docker images.

**Supplementary information:** Supplementary data are available online.

## Introduction

Deep learning (DL) based segmentation methods have become the standard for bioimage analysis, largely surpassing traditional approaches (Jeckel and Drescher, 2021). A bottleneck in the application of DL techniques is the need for comprehensive ground truth (GT) data for training and validation. For the task of cell segmentation, for example, pixel-perfect masks discerning individual cells from the background are crucial. Such data is, however, problem-specific and laborious to produce (Jeckel and Drescher, 2021). Synthesizing data along with its GT is a tried and tested way to alleviate this bottleneck. Cell simulators have been built to form a better understanding of biological growth phenomena, by modeling molecular aspects, rather than aiming at generating photorealistic images (e.g., Gutiérrez *et al*., 2017). For eukaryotic cells, data augmentation using (fluorescence) image generators is well established (e.g., Lehmussola *et al.,* 2007; Svoboda and Ulman, 2017). However, apart from an example for brightfield yeast image generation (Kruitbosch *et al*., 2022), the benefits of synthetic imagine generation have not been exploited for microbial image analysis, in particular in the field of phase-contrast microscopy.

To serve the needs of training data hungry DL-approaches to analyze phase contrast image stacks, we have developed the highly configurable bacterial microcolony simulator *CellSium*. We demonstrate its applicability for object detection (YOLOv5) as well as semantic segmentation (Mask R-CNN) using synthetic data, and verify the resulting neural networks with real image data. Further usage scenarios of *CellSium* beyond image synthetization are also showcased.

## Approach

*CellSium* is an agent-based simulator. Internally, each cell is represented as a Python object, with individual properties, such as shape or growth behavior, implemented as mixins following object-oriented programming paradigms. Cell geometries are modeled via closed polygonal chains, allowing for arbitrary shapes (SI Sec. S.1). Various common ready-to-use unicellular geometries are implemented, such as straight and bent rods, simple coccoid (circular) and ellipsoid shapes (SI Fig. S.2). Colony growth behaviour is implemented in the `grow()` function. Here, phenomenological cell size homeostasis models, such as “timer” or “sizer” can be realized (SI Sec. 2) (Taheri-Araghi *et al*., 2015). The `grow()` method is then called for each simulated time step and cell geometries are used with a physics engine to calculate feasible cell placements. Various output routines exist, such as realistic phase contrast images, time-lapse videos, TrackMate XML (Tinevez *et al*., 2017), or simulation-ready input geometries (STL) for computational fluid dynamics (CFD) simulations (SI Sec. S.3).

The image generation process is shown in Fig. 1A: A “perfect” (noiseless) image is generated (A1-A5), which is deteriorated by adding different kinds of noise. The typical phase contrast “halo” around the cells is generated by Gaussian blurring of the image (A2-4), which is combined to mimic a phase contrast image (A5). Next, uneven illumination is added (A6), along with additive/multiplicative noise (A7, 9), yielding a realistic phase contrast bacterial cell image (A10). All noise models are flexibly configurable. This allows, for instance, simulating deteriorated image quality (SI Sec. S.4), and makes *CellSium* readily transferable to other imaging modalities. Optionally, fluorescence can be generated using a Gaussian point spread function (SI Sec. S.5).

**Figure 1.**
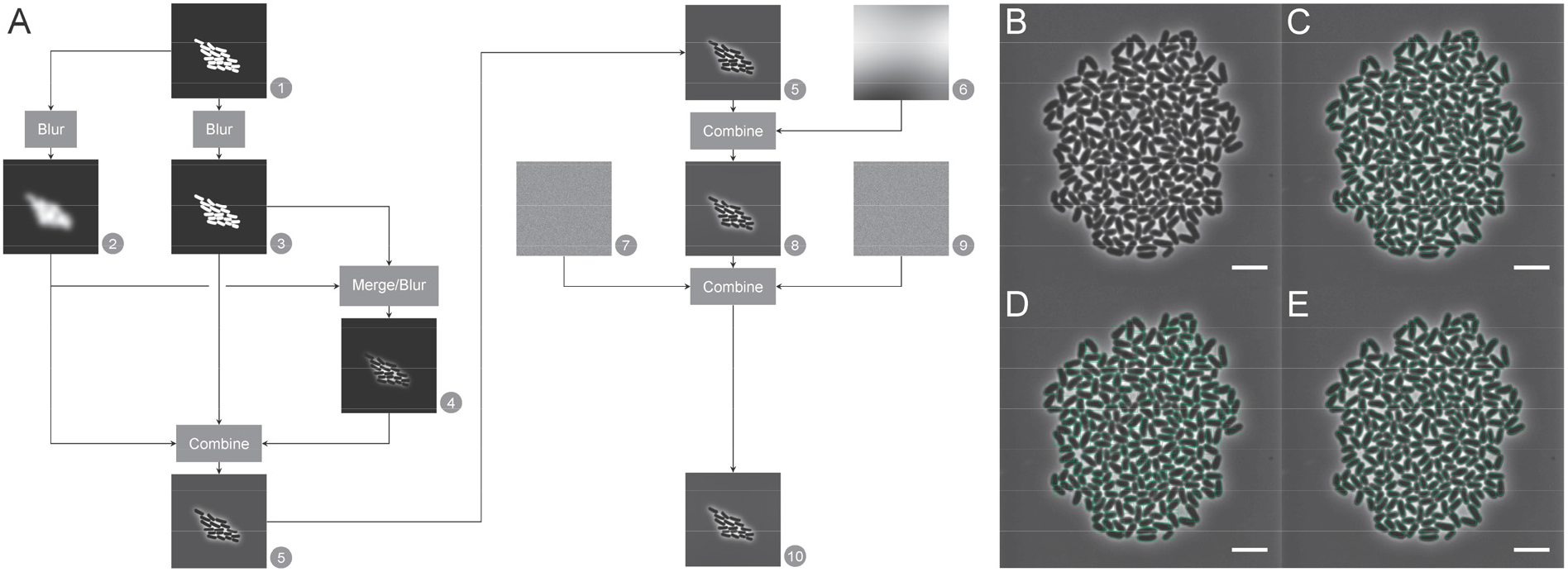
**A**: Highly configurable image generation flow to produce realistic phase contrast images. **B-E**: Prediction results of neural networks trained with data simulated using *CellSium* and evaluated with real data. **B**: input image (a *Corynebacterium glutamicum* microcolony); **C**: manually corrected GT; **D**: YOLOv5 object detector result; **E**: Mask R-CNN segmentation network result.

## Methods

*CellSium* is implemented in the Python programming language, using NumPy and SciPy libraries (github.com/numpy, github.com/scipy/scipy), as well as matplotlib (github.com/matplotlib) and OpenCV (github.com/opencv) for image generation. Physical placement simulation is performed using the off-the-shelve 2D physics library PyMunk (github.com/viblo/pymunk). Source code, PyPI/Anaconda packages, and a Docker image are available, enabling platform-independent usage. The documentation includes API documentation, as well as usage examples for configuring time-lapse simulations, 3D cell geometries, and parametrizable cell models (SI Sec. S.1-3).

### Use case: *CellSium* as GT generator

To assess the applicability of *CellSium* as a GT generator, a data set was generated using the *YOLOOutput* and *COCOOutput* modules (128 images, 512x512, 0.09 μm/pixel, 0 to 512 cells per frame). Synthesized images are verified to have a similar intensity distribution compared to real images with the similar cell/background ratio (SI Sec. S.6). The synthesized outputs were then used to train the object detector/segmentation frameworks YOLOv5 (Jocher et al, 2020) and the Mask R-CNN (He *et al*., 2017) module of MMDetection (Chen *et al*., 2019). As test data, a microcolony image of *Corynebacterium glutamicum* was used (Fig. 1B), which was interactively segmented using the Trainable Weka Segmentation (Arganda-Carreras *et al*., 2017 and then hand-corrected (Fig. 1C).

The YOLOv5 (You Only Look Once v5, <u>ultralytics/yolov5</u> v4.0) net was trained in *yolo5l* mode for 300 epochs, and the test data was predicted with an intersection over union (IoU) threshold for non-maximum suppression (NMS) of 0.6 and a confidence threshold of 0.001. A Mask R-CNN (Region-Convolutional Neural Network) was trained using the MMDetection framework (<u>open-mmlab/mmdetection</u> v2.17.0) for 13 epochs, and the test data was predicted with proposal counts raised to accommodate for the cell count (nms_pre/rpn.max_per_img/rccn.max_per_img 12000/4000/3000 and a score threshold of 0.5).

The mean average precision (mAP) results are given in Table 1. The YOLO net yielded a mAP50 of 0.994, while the Mask R-CNN yielded a mAP_50_ of 0.987 for both bounding boxes segmentations. All files to reproduce the evaluation are available in the GitHub repository github.com/modsim/cellsium.

**Table 1:**
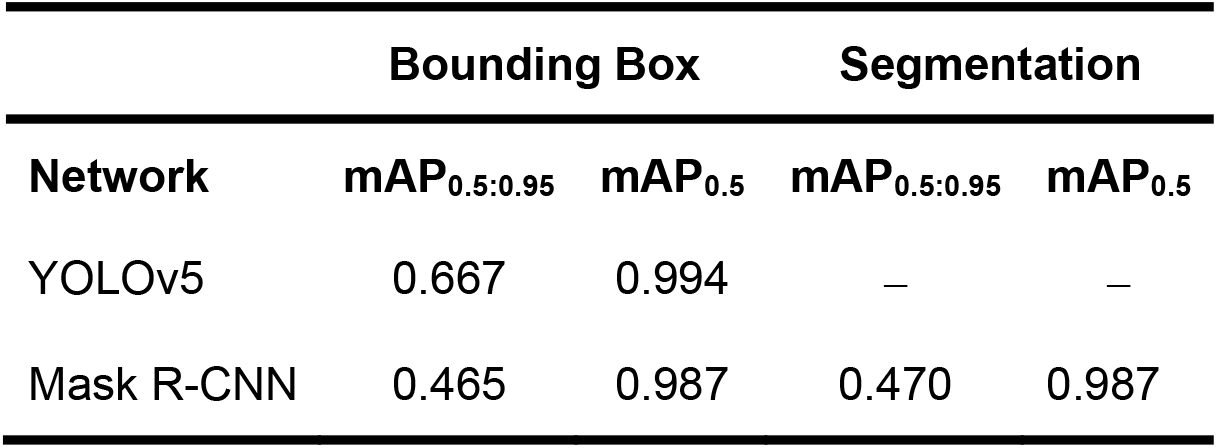
Mean average precision (mAP) results for trained YOLOv5 and Mask R-CNN networks. Tests were performed on an Intel Core i7 4790 (3.6 GHz, 8 threads), 32 GB, GeForce GTX 1080 Ti Linux workstation.

## Conclusion

*CellSium* is a microcolony simulator primarily aimed at bacterial image data set GT generation. Simulated training data showed high intensity histogram correlation with real data and were proven useful for state-of-the-art object detector/segmentation frameworks (YOLOv5, Mask R-CNN), yielding networks capable of producing competitive results with real time-lapse microscopy images. Compared to the laborious manual annotation of image sequences, taking an in silico approach allows generating labelled data with desired characteristics at any amounts, which benefits training and method verification. Likewise, *CellSium* is an effective education tool that allows users, in a well-defined environment, to train method usage or to test phenomenological cell growth models and their impact on the growth characteristics of microcolonies. Finally, developers of bioimage analysis algorithms can benchmark and validate their segmentation and tracking methods with the simulator in flexibly configurable settings.

*CellSium* can be easily used as-is, or, due to its pluggable architecture, serve as a base for implementing other cell types, developing custom cell behavior within growing microcolonies, or embedding the image generation, for example directly in a DL training loop for continuous procedural training data generation. *CellSium*’s current image generation capabilities are well suited for training networks tailored to one imaging modality. A future step, owing to its highly flexible noise model, is to implement generative adversarial networks (GAN) to achieve a high level of realism, independent of the modality.

## Funding

This work was supported by the Deutsche Forschungsgemeinschaft (WI 1705/16-2), the President’s Initiative and Networking Funds of the Helmholtz Association of German Research Centres (SATOMI, ZT-I-PF-04-011), and was performed as part of the Helmholtz School for Data Science in Life, Earth and Energy (HDS-LEE) and received funding from the Helmholtz Association of German Research Centres.

## Notes

### Competing Interest Statement

The authors have declared no competing interest.

### Summary of Updates

Manuscript extended; Supplemental files updated

## References

Arganda-Carreras, I. et al. (2017) Trainable Weka Segmentation: A machine learning tool for microscopy pixel classification. Bioinformatics, 33(15), 2424–2426.

Chen, K. et al. (2019) MMDetection: Open MMLab detection toolbox and benchmark. arXiv:1906.07155.

Gutiérrez, M. et al. (2017) A new improved and extended version of the multicell bacterial simulator gro. ACS Synth. Biol., 6, 1496–1508.

He, K. et al. (2017) Mask R-CNN. In, 2017 IEEE International Conference on Computer Vision (ICCV), pp. 2980–2988.

Jeckel, H. and Drescher, K. (2021) Advances and opportunities in image analysis of bacterial cells and communities. FEMS Microbiol. Rev., 45, 1–14.

Jocher, G. (2020) ultralytics/yolov5: v3.1 - Bug fixes and performance improvements.

Kruitbosch, H.T. et al. (2022) A convolutional neural network for segmentation of yeast cells without manual training annotations. Bioinformatics, 38, 1427–1433.

Lehmussola, A. et al. (2007) Computational framework for simulating fluorescence microscope images with cell populations. IEEE Trans. Med. Imaging, 26, 1010–1016.

Svoboda, D. and Ulman, V. (2017) MitoGen: A framework for generating 3D synthetic timelapse sequences of cell populations in fluorescence microscopy. IEEE Trans. Med. Imaging, 36, 310–321.

Taheri-Araghi, S. et al. (2015). Cell-size control and homeostasis in bacteria. Current Biology, 25(3), 385–391.

Tinevez, J.-Y. et al. (2017) TrackMate: An open and extensible platform for single-particle tracking. Methods, 115, 80–90.

