## Supplementary Information for "*CellSium* – Versatile Cell Simulator for Microcolony Ground Truth Generation"

### S.1 Cell Shape Geometries

*CellSium* allows to customize the individual cell shape, its appearance and evolution in time. Shapes are modelled using non-overlapping closed polygon shape in 2D, cells are rendered and placed conforming with physics without overlap/collision. This way, the set of simulated morphologies can be easily extended and dynamically adapted over time. To demonstrate these capabilities we simulated an artificial growing tree colony (see Fig. S.1).

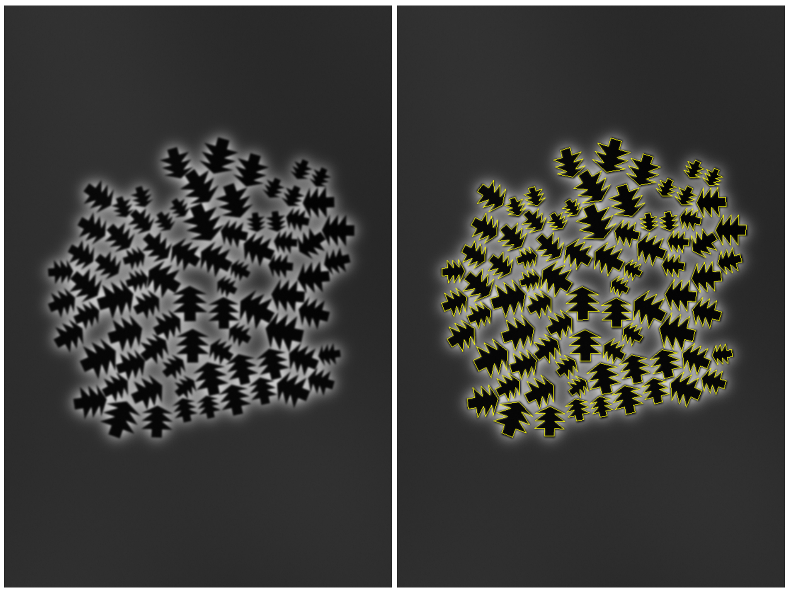

Fig. S.1: Artificial tree colonies without (left) and with (right) contours highlighted.

*CellSium* comes with several predefined microbial shapes that are commonly studied with microfluidic live-cell imaging. Figure S.2 shows these ready-to-use bacterial cell geometries implemented in *CellSium*.

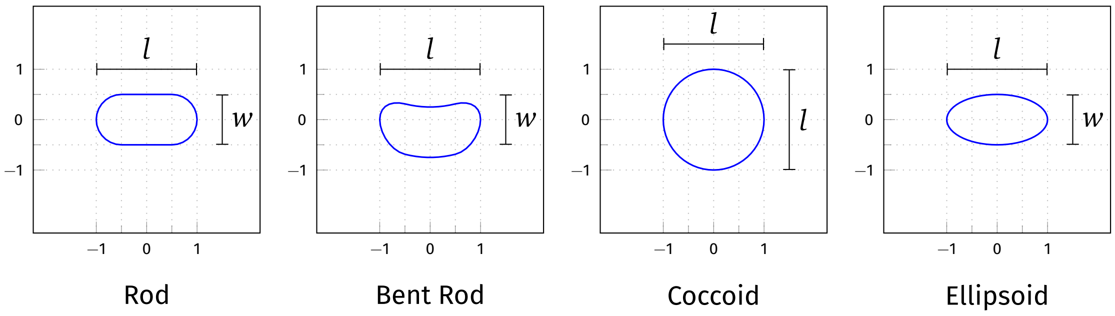


Fig. S.2: Implemented procedurally generated shapes. All shapes shown for $l = 2$, $w = 1$ [a.u.].

### S.2 Usage Example: Simulation of Time-lapse Sequences to Visualize and Analyze Cellular Growth Behaviour

How cells keep a stable size over time, despite various stochasticity in the underlying cellular processes, is a question that fascinates cell biologists since decades. Here, we show how *CellSium* can be used to produce synthetic time-lapse image sequences that render prominent cellular size models and derive characteristic distributions, such as for the cell length. This is exemplified for the so-called "Timer" and the "Sizer" models (Taheri-Araghi *et al.*, 2015). The "Timer" model assumed fixed time intervals between division events, whereas the "Sizer" model prescribes cell division after a specific cell size is reached. Python encodings for both models is given in listings 1 and 2.

```python
class TimerCell(SimulatedCell):
    @staticmethod
    def random_sequences(sequence: Any) -> Mapping[str, Any]:
        return dict(elongation_rate=sequence.normal(1.5, 0.25))  # µm·h^-1

    def birth(
        self, parent: Optional["TimerCell"] = None, ts: Optional[Timestep] = None
    ) -> None:
        self.elongation_rate = next(self.random.elongation_rate)
        self.division_time = h_to_s(1.0)

    def grow(self, ts: Timestep) -> None:
        self.length += self.elongation_rate * ts.hours

        if ts.time > (self.birth_time + self.division_time):
            offspring_a, offspring_b = self.divide(ts)
            offspring_a.length = offspring_b.length = self.length / 2
```

Listing 1: `TimerCell` class implementing a simple “Timer” model.

```python
class SizerCell(SimulatedCell):
    @staticmethod
    def random_sequences(sequence: Any) -> Mapping[str, Any]:
        return dict(division_size=sequence.normal(3.0, 0.25))  # µm

    def birth(
        self, parent: Optional["SizerCell"] = None, ts: Optional[Timestep] = None
    ) -> None:
        self.division_size = next(self.random.division_size)
        self.elongation_rate = 1.5

    def grow(self, ts: Timestep) -> None:
        self.length += self.elongation_rate * ts.hours

        if self.length > self.division_size:
            offspring_a, offspring_b = self.divide(ts)
            offspring_a.length = offspring_b.length = self.length / 2
```

Listing 2: `SizerCell` class implementing a simple “Sizer” model.

In Fig. S.3 the results of a simulation ($0 \leq t \leq 12\,\mathrm{h}$) with the two vanilla "Timer" and "Sizer" models are shown, as given in the listings. Both models produce exponential growth with a growth rate of approximately $0.68\,\mathrm{h}^{-1}$ ($R^2 > 0.99$). As long as no randomness is added apart from the elongation rate, for the “Timer” model all cells double in a synchronous fashion, resulting in step-wise increasing cell numbers. Here, the “Sizer” model produces a smoother growth curve. Comparing the cell length distributions derived from the simulations with the two models in Fig. S.3 (bottom row) shows that despite the differences in distributions, both simulations result in very similar average cell lengths of approximately $2.15\,\mathrm{\mu m}$. Furthermore, it is easily possible to add various stochastic effects to the vanilla models. This may be useful for inferring measured distributions.

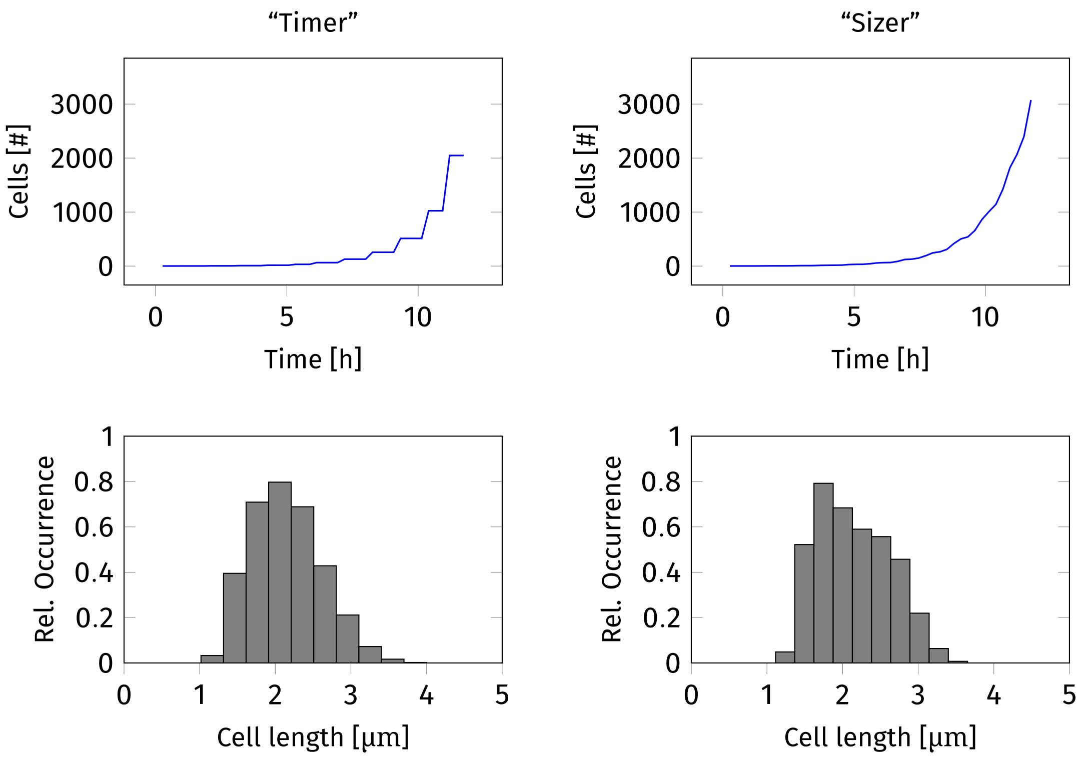


Fig. S.3: Example growth curves derived from simulation experiments. Starting with a single, randomized cell, the "Timer" and the "Sizer" models were simulated with *CellSium*. Simulation snapshots were taken for 12 h in 0.25 h intervals. The “Timer” model yielded a growth rate of 0.681 $\mathrm{h}^{-1}$, the “Sizer” of 0.685 $\mathrm{h}^{-1}$. In the bottom row, corresponding cell length distributions are shown. For the “Timer” example a mean cell length of 2.15±0.46 μm over 13 275 cells was observed, whereas for the “Sizer” the average length was 2.15±0.48 μm over 17 862 cells.

### S.3 Usage Example: Cell and Microcolony Geometries for Computational Fluid Dynamics Simulations

Computational fluid dynamics (CFD) studies provide the ability to shed light on the impact physical conditions, such as nutrient concentrations or flow velocity, have on growing cells and microcolonies, see for example Westerwalbesloh *et al.* (2015). Here, we show how *CellSium* is used to automatically create input geometries of a bacterial microcolony from data acquired by live-cell imaging and image analysis. To this end, the .stl-output is used, producing cells as 3D solids-of-revolution, based on rotating their 2D representation. An example is given in Fig. S.4. Taken the 3D geometry of the microcolony, we produced a mesh using COMSOL® Multiphysics (ver. 5.3) and simulated glucose diffusion. To this end, the cells were placed within a cube of 40 µm × 60 µm × 1.2 µm "filled" with medium. A constant glucose concentration of 222 mol m$^{-3}$ was specified at the top/bottom surfaces of the cube (i.e. at y min/max in Fig. S.5), to model substrate replenishment via the nutrient channels. A more detailed description of microfluidic chip design is given in Westerwalbesloh *et al.* (2015). The simulated glucose concentration distribution is visualized in Fig. S.5.

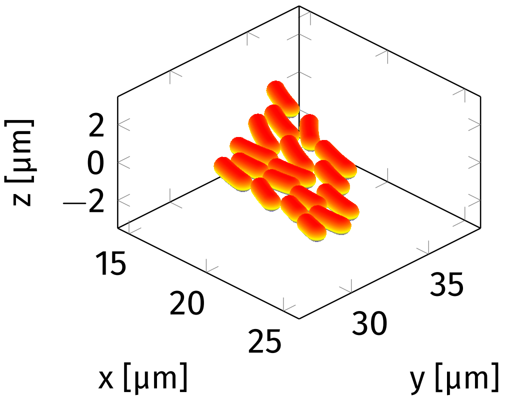


(a) Mesh Overview

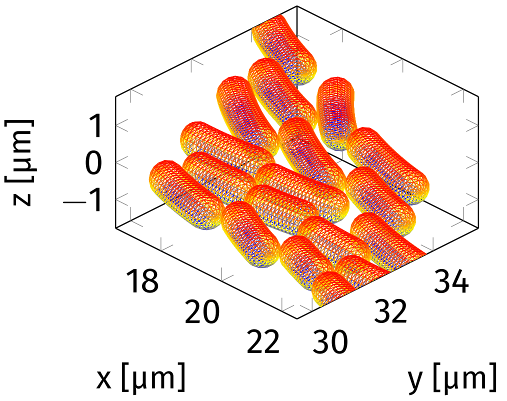


(b) Mesh close-up

Fig. S.4: Mesh examples generated using *CellSium*.

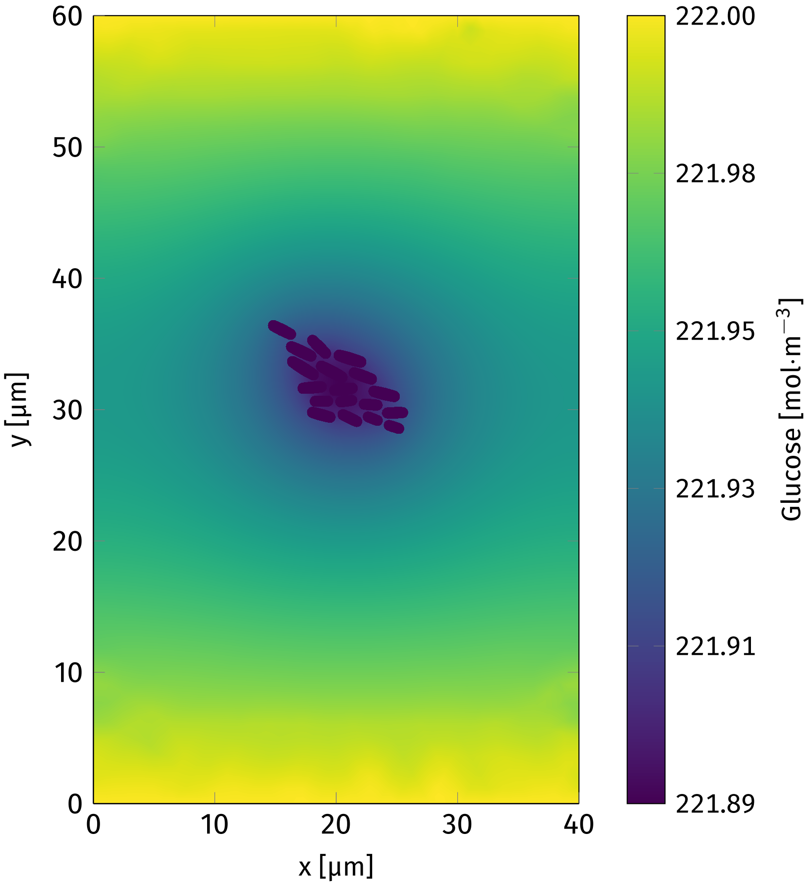


Fig. S.5: Glucose distribution at steady-state in a rectangular growth chamber, simulated result produced using COMSOL® Multiphysics. Cells were assumed to take up glucose at a rate of $1.14 \cdot 10^{-6}$ mol s$^{-1}$ m$^{-2}$, with a glucose diffusion coefficient of $5.4 \cdot 10^{-10}$ m$^2$ s$^{-1}$ at 30 °C (Westerwalbesloh *et al.*, 2015). The mesh was generated using COMSOL® with the 'extra fine' pre-set for general physics, yielding a total of 114 843 elements.

### S.4 Usage Example: Simulating Imaging Challenges - Focus Loss

Imaging microbial development is a challenging process and several difficulties occur in practice. For example, low-contrast to noise ratio, illumination changes and focus loss due to temperature changes or mechanical movement are common in microfluidic live-cell imaging. Mimicking these phenomena in the *CellSium* simulator is possible, by applying custom modifications to the generated images. As an example, the result of a simulated focus loss with different degrees of blurring is shown in Fig. S.6. Such images along with their ground truth help to diversify the training data set.

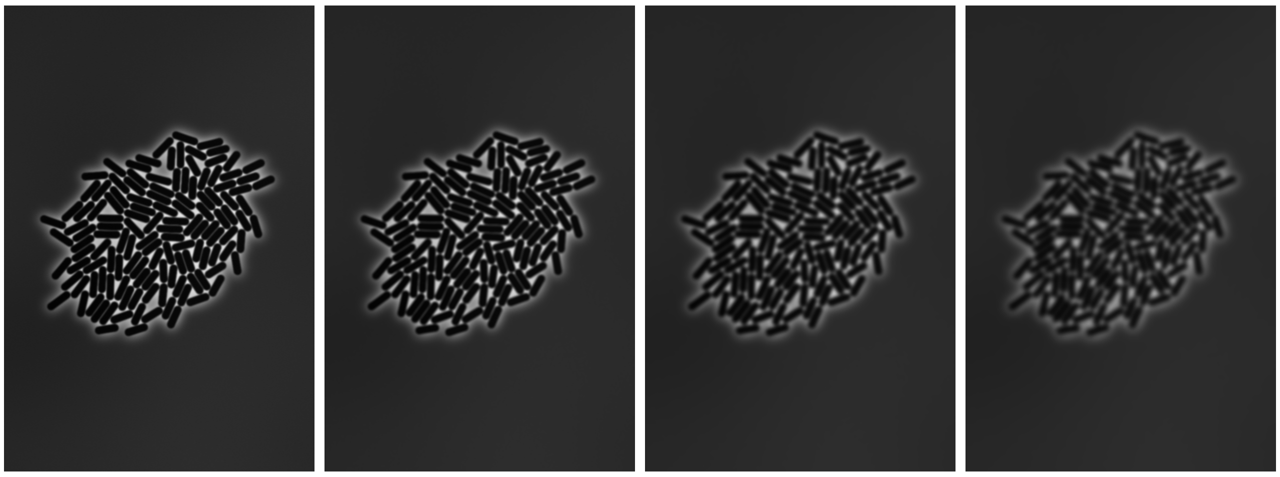

(a) Original (b) 1x (c) 5x (d) 10x

Fig. S.6: Simulating focus loss. A normalized box filter (5 × 5 kernel) is applied 1, 5 and 10 times to the original image generated with CellSium to simulate various degrees of out-of-focus images.

### S.5 *Cellsium* Feature: Fluorescence Image Generation

To create synthetic fluorescence images with *CellSium*, the values of a simplified point spread function (PSF) evaluation are repeatedly drawn onto an empty canvas. In the implementation, random coordinates within each cell are picked and used as emitter positions. As PSF, a Gaussian function is used, being added to the image canvas for every emitter position. Fluorescence intensity is modeled by scattering more or less emitters into the cells, normalized by cell area and dependent on the fluorescence parameter of the cell. This approach models fluorophores which are equally distributed within the bacterial cytosol, an example with two differing levels of fluorescence intensity is shown in Fig. S.7.

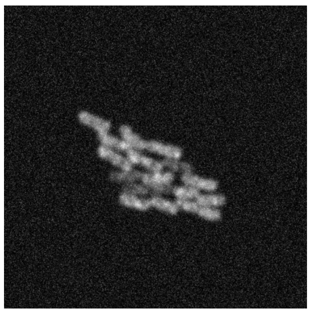

Fig. S.7: Synthetic fluorescence image with randomly assigned fluorescences (two different levels), generated with *CellSium*. Fluorescence is simulated by randomly scattering point spread functions in the cell area.

### S.6 Similarity of Real and Synthesized Images

A quantitative comparison of the similarity between real microscopic images and the synthetic images, generated with *CellSium*, is based on comparing their corresponding intensity histograms. For that, we pick crops of the images, containing similar amounts of cells and cell/background ratios. Figure S.8 shows the resulting histograms for the synthetic and real images, and their correlation. For the histograms, we can obtain 75% Pearson correlation coefficient (PCC), indicating high similarity.

The PCC of the histogram pairs can be used as a metric to optimize the parameter of *Cellsium* to support other microscopic modalities (e.g., bright-field imaging), or to develop tailored noise models.

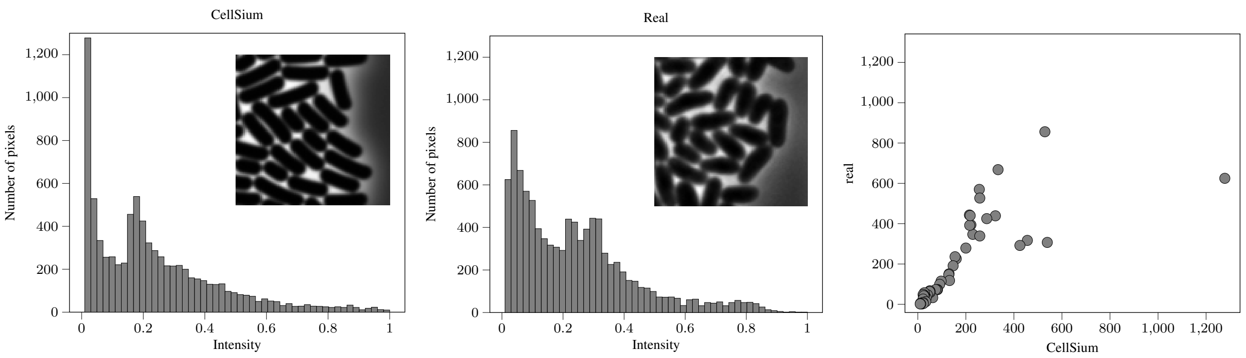


(a) (b) (c)

Fig. S.8: The histograms (50 bins) of the normalized synthetic S.8a and real S.8b images and their correlation plot Fig. S.8c

### References

Taheri-Araghi, S., Bradde, S., Sauls, J. T., Hill, N. S., Levin, P. A., Paulsson, J., Vergassola, M., and Jun, S. (2015). Cell-size control and homeostasis in bacteria. *Current Biology*, **25**(3), 385–391.

Westerwalbesloh, C., Grünberger, A., Stute, B., Weber, S., Wiechert, W., Kohlheyer, D., and von Lieres, E. (2015). Modeling and CFD simulation of nutrient distribution in picoliter bioreactors for bacterial growth studies on single-cell level. *Lab on a Chip*, **15**, 4177–4186.
